# Possible Implications of AlphaFold2 for Crystallographic Phasing by Molecular Replacement

**DOI:** 10.1101/2021.05.18.444614

**Authors:** Airlie J. McCoy, Massimo D. Sammito, Randy J. Read

## Abstract

The AlphaFold2 results in the 14^th^ edition of Critical Assessment of Structure Prediction (CASP14) showed that accurate (low root-mean-square deviation) *in silico* models of protein structure domains are on the horizon, whether or not the protein is related to known structures through high-coverage sequence similarity. As highly accurate models become available, generated by harnessing the power of correlated mutations and deep learning, one of the aspects of structural biology to be impacted will be methods of phasing in crystallography. We here use the data from CASP14 to explore the prospect for changes in phasing methods, and in particular to explore the prospects for molecular replacement phasing using *in silico* models.

**Synopsis:** We discuss the implications of the AlphaFold2 protein structure modelling software for crystallographic phasing strategies.

## 1. Introduction

The quality of a model for phasing a crystal structure by molecular replacement, for a given diffraction resolution limit, depends on the model’s root-mean-square deviation (*rmsd*) to the target structure and fraction of the total scattering (*f_m_*) it represents. Broadly, as the resolution of the experimental data decreases, the *f_m_* must increase (Figure 1). Specific predictions based on *rmsd, f_m_* and data resolution guide molecular replacement phasing strategies in individual cases (Oeffner *et al*., 2018)

**Figure 1.**
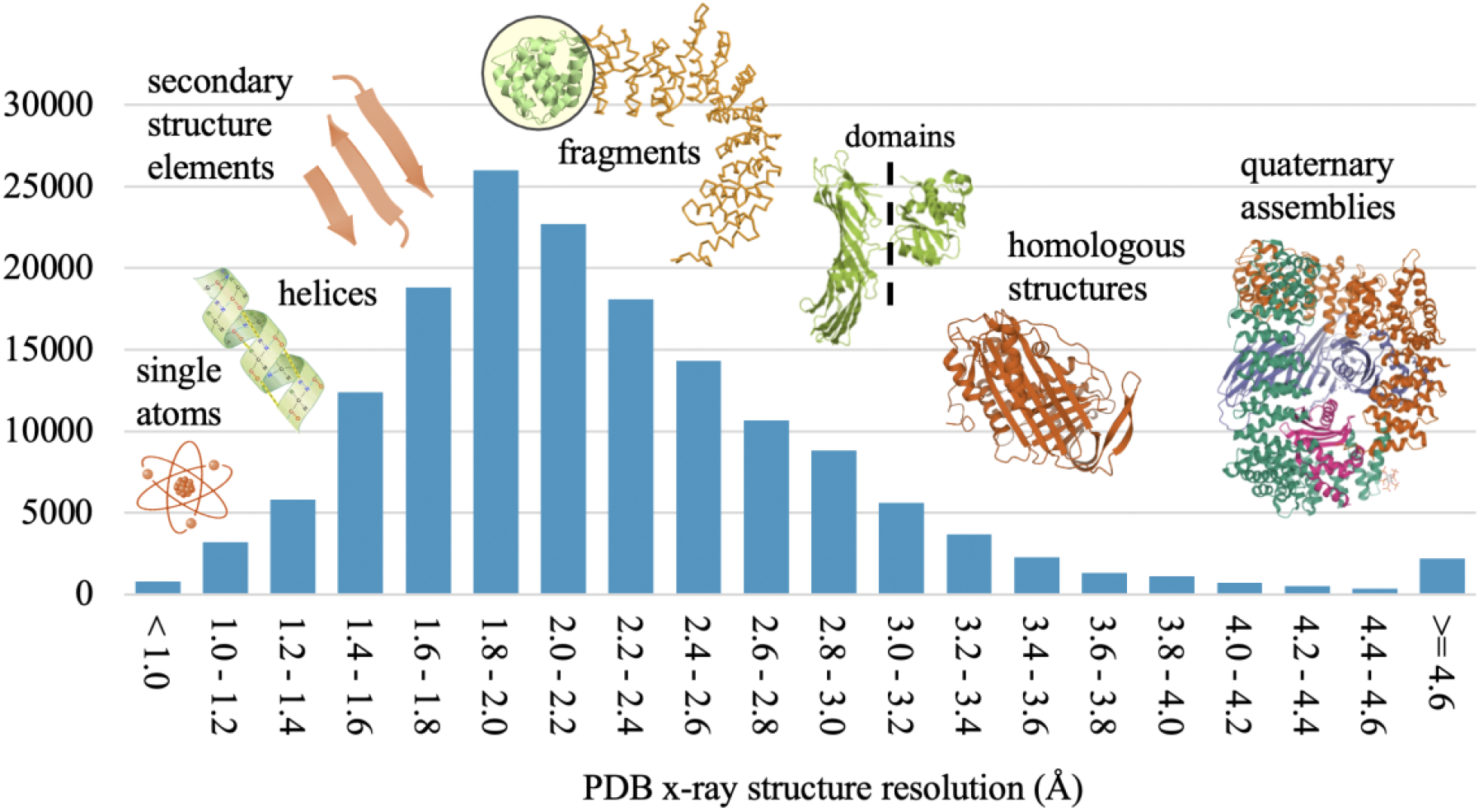
Histogram of the distribution of structures in the PDB by resolution. The relationship between resolution of the data and the size of the models that are appropriate for molecular replacement is indicated.

Where data resolution extends to better than ~1 Å, the model can be as tiny as a single atom. A single atom can be considered the perfect *in silico* substructure model, having an *rmsd* of zero to the target structure, although, overall, the structure factors calculated from this model have large errors because of the extremely low *f_m_*. Following single atom molecular replacement, log-likelihood gradient completion can rapidly locate the remaining ordered atoms and for these resolutions the phase problem is considered solved (McCoy *et al*., 2017).

Small secondary structure elements of helix or beta-sheet are viable models when data extend to better than ~2.2 Å resolution, and in some cases this technique extends to resolutions of 2.5 Å. The model’s atomic coordinates can be extracted from known structures or generated *in silico* (Glykos & Kokkinidis, 2001). Success does not necessarily require a homologous model with a sequence identity over 30%, despite this being a commonly quoted metric for molecular replacement success (Scapin, 2013). Density modification and model building are central to structure completion with this method. Accurate fragments are regularly used for molecular replacement in software such as *Arcimboldo* (Sammito *et al*., 2015, 2014; Rodríguez *et al*., 2009) and *Ample* (Bibby *et al*., 2012; Rigden *et al*., 2018; Simpkin *et al*., 2019). For these resolutions, the phase problem is also largely considered solved. The median resolution in the PDB is 2.2 Å, making this approach possible for many present-day crystallographic structures.

Where the experimental data extend to lower than ~2.2 Å resolution the models required for molecular replacement must represent, to at least some degree, the fold of the target protein (the hydrophobic core or more). A high *f_m_* and a low *rmsd* become progressively more important as the resolution decreases and by ~3.0 Å resolution, in a typical crystal, a whole-structure model with less than 1 Å *rmsd* would be required for successful molecular replacement and model completion. This is the zone (sub-3.0 Å) in which homologs, template-based modelling and *in silico* models become particularly valuable.

For those targets distantly related to a homologous structure, early attempts at template-based modelling (as catalogued by the Critical Assessment of Structure Prediction, CASP) generally increased (rather than decreased, as was the aim) the *rmsd* to the target. CASP7 was the first CASP to show that template-based modelling improved models for molecular replacement. From this CASP also came the first case in which an *in silico* structure prediction for a natural protein with an asymmetric, globular fold was successfully used for molecular replacement, albeit retrospectively (Qian *et al*., 2007). Since then, an *in silico* model has been used in the real-world molecular replacement phasing of peptidoglycan polymerase RodA (Sjodt *et al*., 2018).

Since CASP7, CASP has included a metric for scoring individual model predictions based on their usefulness in molecular replacement, with steady progress each challenge round. From CASP13 it was demonstrated that not only accurate coordinates, but also accurate estimates of the errors in the coordinates were critical for successful molecular replacement (Croll *et al*., 2019).

There are several pipelines for molecular replacement with *in silico* models. One of the first was CaspR, making use of models produced by MODELLER (Claude et al., 2004). The first iteration of the Ample pipeline developed a cluster-and-truncate approach to the use of rapidly computed *ab initio* models generated by Rosetta or QUARK (Bibby *et al*., 2012). In further developments, Ample has been extended to use structure predictions from the GREMLIN and PconsFam databases (Simpkin *et al*., 2019). Models from I-TASSER, generated by full-length iterative structural fragment reassembly, have been incorporated in the I-TASSER-MR server, which uses progressive sequence truncation to edit the models for molecular replacement (Wang *et al*., 2017). AWSEM-Suite combines both homology model templates and coevolutionary information with the physico-chemical energy terms of AWSEM (Jin *et al*., 2020). In our own collaborations, the *phenix.mr_rosetta* (Terwilliger *et al*., 2012) pipeline can use Rosetta to rebuild template structures prior to attempting molecular replacement. We also use *Rosetta*, extended to includes a term for fit to the electron density (DiMaio, 2013) to rank putative molecular replacement solutions, and to rebuild very poor models after molecular replacement.

## 2. CASP14

CASP14 has established a leap in protein structure prediction. The primary CASP metric for ranking models and modelling groups is the GDT_TS (Global Distance Test Total Score), a structure similarity measure designed and developed for the structure alignment program LGA (Local-Global Alignment) as an alternative to the *rmsd* (Zemla, 2003). The GDT measures the percentage of C*a* that are found within certain distance cut-offs of one another between model and target (either dependent upon or independent of a sequence alignment): the cut-off distance(s) must be defined for each reported GDT value. The GDT_TS is the average of four cut-off distances (1, 2, 4, and 8 Å). Higher GDT values are achieved with better models, in contrast to the *rmsd* where lower values are better.

Harnessing the power of correlated mutations, contact predictions and deep learning, the AlphaFold2 group (group 427 in the CASP14 numbering) from the commercial organisation DeepMind (Service, 2018; Callaway, 2020) were ranked first on Z-score for GDT_TS, reaching values over twice, and up to thrice, that of the second and subsequent ranked groups, depending on classification of targets considered. This was achieved in the background of major improvements from other groups, including the Baker group (BAKER group 473, ranked second and BAKER-experimental group 403, ranked third) (Hiranuma *et al*., 2021), who used similar methods in academic settings.

For the first time, structures submitted to CASP as targets were phased with the help of models provided during the assessment. For target T1058, the structure was solved by MR-SAD with the AlphaFold2 models after attempting molecular replacement with homologous structures, domains thereof, and server models (Tidow, 2020). For T1089, the AlphaFold2 models gave a far higher molecular replacement signal than using trimmed ensemble models (Rees, 2020). For T1100, several of the models submitted to CASP, including AlphaFold2, gave a molecular replacement solution, where NMR structures of individual domains failed (Lupas *et al*., 2020). The AlphaFold2 model for T1064 has also been used to solve Sars-Cov-2 ORF8 retrospectively by molecular replacement (Flower & Hurley, 2021).

## 3. Molecular replacement assessment

The suitability for molecular replacement is one of the metrics for high accuracy assessment in CASP including CASP14. The assessment uses a log-likelihood gain (LLG) calculated in *Phaser* (McCoy *et al*., 2007; Read & McCoy, 2016). The likelihood is the probability that the data would have been measured, given the model, and the log-likelihood gain is the difference between the log-likelihood of the model and that calculated from a random distribution of the same atoms (Wilson, 1949).

An important component of the scoring of models with the LLG is the incorporation of estimated error in coordinates. As part of the modelling, groups are encouraged to estimate the error (Δ) in each atomic position and to record that estimate in the B-factor column of the deposited PDB file. This error estimate can be converted to a B-factor for each atomic position through the relationship *B*=8*π*^2^Δ^2^/3 and thereby used to weight each atom in the LLG calculation appropriately. Accurate estimates of the error improve the LLG (Bunkóczi *et al*., 2015; Croll *et al*., 2019), and in practice, will add value to the models by increasing the signal in molecular replacement searches.

If, during molecular replacement searches, a pose of a model has an LLG over a certain space-group dependent value (60 in non-polar space groups; 50 in polar space groups; and 30 in P1), the pose is probably correct. However, achieving this LLG is not sufficient to determine whether the full structure can be traced and refined to a point suitable for interpretation, publication and deposition; this also depends on the accuracy and completeness of the model, and the resolution of the data. Testing the ability of a model to phase includes validation of these steps downstream of nominally successful model replacement.

### 3.1. Model parameters

Before structure solution, the LLG that will be achieved for a correct pose of a model can be estimated with the ‘expected LLG’ (eLLG; (McCoy *et al*., 2017)). We have previously shown that the eLLG for each reflection *hkl* can be calculated from the σ_A_ parameter (Read, 1986).

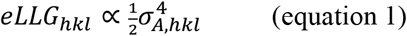

The total eLLG is the sum of this value over all reflections *hkl*. The resolution-dependent *σ_A_* term is approximated with a four-parameter curve which includes the *rmsd*, the *f_m_* and also two solvent parameters which affect the σ_A_ value at resolutions lower than approximately 8 Å (Murshudov *et al*., 1997). For reflections higher than 8 Å resolution, this curve is dominated by the dependence on the square-root of*f_m_* and an exponential dependence on *rmsd*^2^. For each reflection *hkl* (resolution *d_hkl_*),

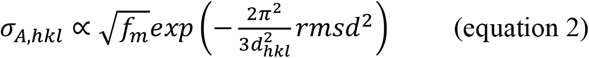

Since the eLLG is a good estimate of the LLG when there are no pathologies in the data (Oeffner *et al*., 2018), the above equation also shows the relationship between *rmsd, f_m_* and the LLG. The dependence of the LLG on *rmsd* and *f_m_* is shown in Figure 2.

**Figure 2.**
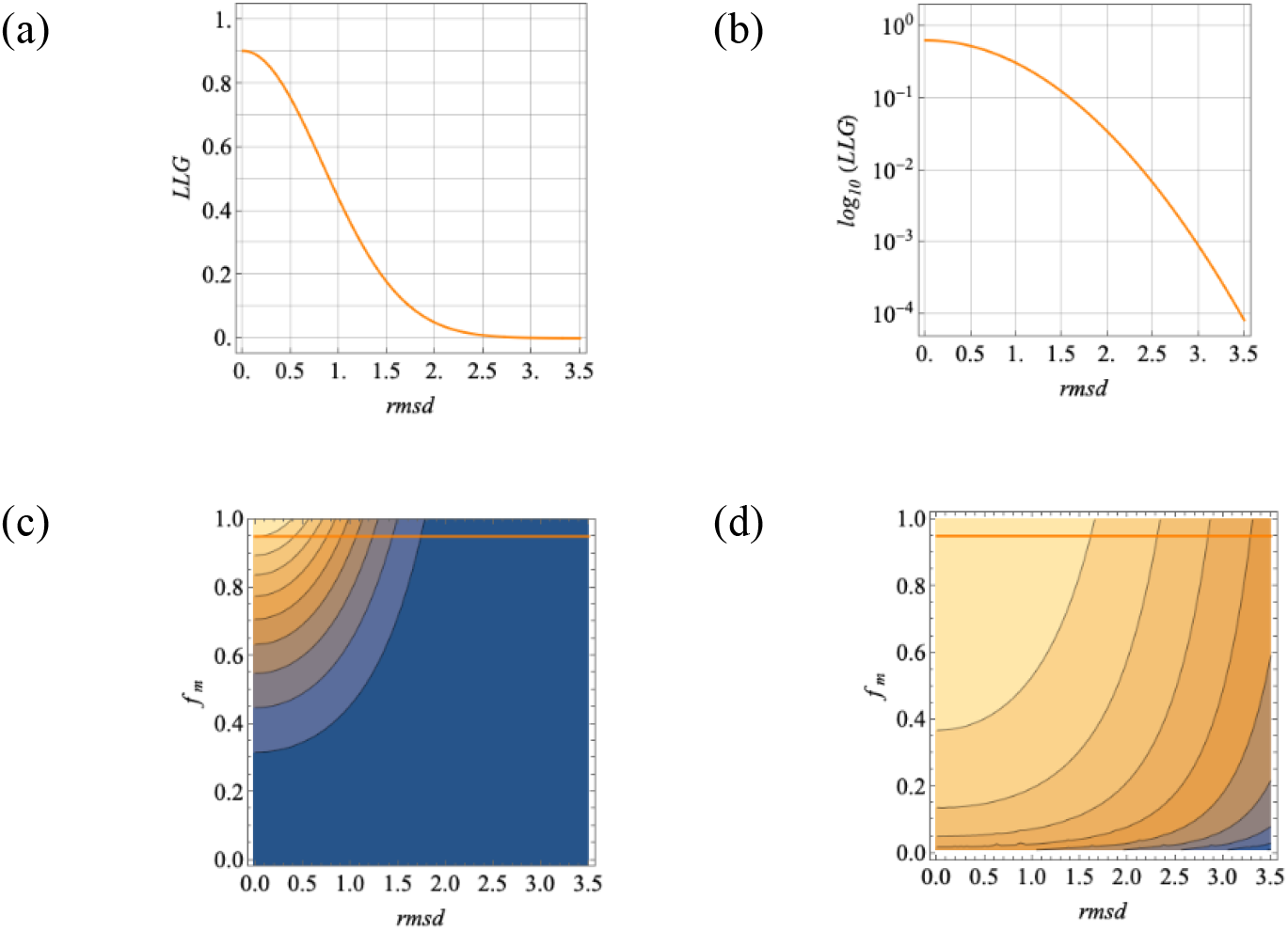
Illustration of the relationship between *rmsd, f_m_* and LLG per reflection (equation 2). (a) LLG for *f_m_* = 0.8 linear scale (b) LLG for *f_m_* = 0.8 logarithmic scale (c) contour plot showing LLG for *rmsd* versus *f_m_*, linear scale (d) contour plot showing for *rmsd* versus *f_m_*, logarithmic scale. The value *f_m_* = 0.8 is shown with an orange line on the contour plots.

Any parameter that correlates strongly with *rmsd* (such as the GDT_HA, when the GDT_HA is high) will show the same relationship to the LLG.

The *rmsd* is only useful for predicting the LLG when the same *rmsd* value describes the difference between the model and target both locally and globally. If it varies between regions, the *rmsd* is dominated by the regions where the *rmsd* is large, whereas the LLG score will be dominated by the regions where the *rmsd* is low. In practice, there will be regions that are better modelled than others, and the LLG obtained from a model will be higher when the estimated errors in each coordinate are good and are incorporated into the electron density calculation via the B-factor, as described above.

### 3.2. Targets

There were 33 crystal structures provided as modelling targets for CASP14. Of these, 31 have a single protein sequence (sometimes in multiple copies) in the asymmetric unit, and two crystals have two protein sequences. The relationship between the naming of the targets and the crystal structures is not straightforward. In 30 of 31 cases with a single sequence, there is a one-to-one correspondence between a sequence and a CASP target number (e.g. T1032). The exception is the case of crystal 6vr4, where there are 9 separate targets (T1031 T1033 T1035 T1037 T1039 T1040 T1041 T1042 T1043), each representing between 95 and 404 residues of the full polypeptide sequence of 2194 residues, two copies of which are present in the asymmetric unit. In 11 of 30 cases the full sequence is deemed to be ‘multidom’ (multi-domain) and is also divided into two, three or four domains and these treated as additional, separate targets, with the suffix “-D1”, “-D2”, “-D3”, or “-D4” added to the target number for the whole structure (e.g. T1024 divided into T1024-D1 and T1024-D2). In 8 of 11 ‘multidom’ targets, the whole sequence target is referenced suffixed with “-D0” (e.g. T1038-D0) and not simply by the target number alone, as in the other 3 of 11 (e.g. T1024). The 19 of 30 that are not ‘multidom’ have a single domain defined within the full sequence, and the target is given the suffix “-D1” (e.g. T1032-D1). In the two of 33 where there are two sequences in the asymmetric unit, the corresponding two targets are named with the same target number with the suffix “s1” and “s2” added (T1046s1 and T1046s2; T1065s1 and T1065s2). Neither of these divide their constituents into domains, and the single targets are given the “-D1” suffix.

We also considered one other crystal, 6un9, corresponding to target T1048, which was cancelled from CASP14 for lack of tertiary structure; it is a single sequence that folds into a single helix and forms a coiled-coil. A model for this structure was also prepared by the AlphaFold2 group before it was cancelled.

In total, we considered 72 CASP14 targets from the 34 crystal structures, when the domains of ‘multidom’ targets are included in the total (Table 1).

**Table 1.** The 34 crystal structures included in CASP14 and the targets associated with each crystal.

| Crystal Number | Target Number | CASP Target | Residues | CASP Domain | Residues | 'Multidom' Domains(s) | Residues |
| --- | --- | --- | --- | --- | --- | --- | --- |
| 1 | 1 | T1024 | 408 |  |  | T1024-D1 | 193 |
|  |  |  |  |  |  | T1024-D2 | 204 |
| 2 | 2 | T1030 | 273 | T1030-D0 | (273) | T1030-D1 | 154 |
|  |  |  |  |  |  | T1030-D2 | 119 |
| 3 | 3 | T1031 | 95 | T1031-D1 | (95) |  |  |
|  | 4 | T1033 | 100 | T1033-D1 | (100) |  |  |
|  | 5 | T1035 | 102 | T1035-D1 | (102) |  |  |
|  | 6 | T1037 | 404 | T1037-D1 | (404) |  |  |
|  | 7 | T1039 | 161 | T1039-D1 | (161) |  |  |
|  | 8 | T1040 | 130 | T1040-D1 | (130) |  |  |
|  | 9 | T1041 | 242 | T1041-D1 | (242) |  |  |
|  | 10 | T1042 | 289 | T1042-D1 | 276 |  |  |
|  | 11 | T1043 | 148 | T1043-D1 | (148) |  |  |
| 4 | 12 | T1032 | 284 | T1032-D1 | 169 |  |  |
| 5 | 13 | T1034 | 156 | T1034-D1 | (156) |  |  |
| 6 | 14 | T1038 | 199 | T1038-D0 | 190 | T1038-D1 | 114 |
|  |  |  |  |  |  | T1038-D2 | 76 |
| 7 | 15 | T1046s1 | 216 | T1046s1-D1 | 72 |  |  |
|  | 16 | T1046s2 |  | T1046s2-D1 | 141 |  |  |
| 8 | 17 | T1048* |  |  |  |  |  |
| 9 | 18 | T1049 | 141 | T1049-D1 | 134 |  |  |
| 10 | 19 | T1050 | 779 |  |  | T1050-D1 | 321 |
|  |  |  |  |  |  | T1050-D2 | 316 |
|  |  |  |  |  |  | T1050-D3 | 128 |
| 11 | 20 | T1052 | 832 | T1052-D0 | (832) |  |  |
| 12 | 21 | T1053 | 580 | T1053-D0 | 576 | T1053-D1 | 405 |
|  |  |  |  |  |  | T1053-D2 | 171 |
| 13 | 22 | T1054 | 190 | T1054-D1 | 143 |  |  |
| 14 | 23 | T1056 | 186 | T1056-D1 | 169 |  |  |
| 15 | 24 | T1058 | 382 | T1058-D0 | (382) | T1058-D1 | 221 |
|  |  |  |  |  |  | T1058-D2 | 161 |
| 16 | 25 | T1064 | 106 | T1064-D1 | 92 |  |  |
| 17 | 26 | T1065s1 | 225 | T1065s1-D1 | 119 |  |  |
|  | 27 | T1065s2 |  | T1065s2-D1 | 98 |  |  |
| 18 | 28 | T1067 | 220 | T1067-D1 | (221) |  |  |
| 19 | 29 | T1070 | 335 |  |  | T1070-D1 | 76 |
|  |  |  |  |  |  | T1070-D2 | 101 |
|  |  |  |  |  |  | T1070-D3 | 76 |
|  |  |  |  |  |  | T1070-D4 | 68 |
| 20 | 30 | T1073 | 58 | T1073-D1 | (59) |  |  |
| 21 | 31 | T1074 | 131 | T1074-D1 | (132) |  |  |
| 22 | 32 | T1079 | 483 | T1079-D1 | 451 |  |  |
| 23 | 33 | T1080 | 137 | T1080-D1 | 133 |  |  |
| 24 | 34 | T1082 | 97 | T1082-D1 | 75 |  |  |
| 25 | 35 | T1083 | 196 | T1083-D1 | 92 |  |  |
| 26 | 36 | T1084 | 146 | T1084-D1 | 71 |  |  |
| 27 | 37 | T1085 | 588 | T1085-D0 | 406 | T1085-D1 | 167 |
|  |  |  |  |  |  | T1085-D2 | 182 |
|  |  |  |  |  |  | T1085-D3 | 57 |
| 28 | 38 | T1086 | 408 | T1086-D0 | 381 | T1086-D1 | 193 |
|  |  |  |  |  |  | T1086-D2 | 188 |
| 29 | 39 | T1087 | 186 | T1087-D1 | 93 |  |  |
| 30 | 40 | T1089 | 404 | T1089-D1 | 377 |  |  |
| 31 | 41 | T1090 | 193 | T1090-D1 | 191 |  |  |
| 32 | 42 | T1091 | 863 |  |  | T1091-D1 | 139 |
|  |  |  |  |  |  | T1091-D2 | 107 |
|  |  |  |  |  |  | T1091-D3 | 106 |
|  |  |  |  |  |  | T1091-D4 | 112 |
| 33 | 43 | T1100 | 338 |  |  | T1100-D1 | 171 |
|  |  |  |  |  |  | T1100-D2 | 166 |
| 34 | 44 | T1101 | 318 | T1101-D0 | 307 | T1101-D1 | 83 |
|  |  |  |  |  |  | T1101-D2 | 224 |

\* cancelled

### 3.3. Classification of targets

CASP classifies the targets by modelling difficulty in four categories: free modelling (FM), template-based modelling (TBM-easy and TBM-hard), and structures on the boundary between free modelling and template-based modelling (FM/TBM). In the set of crystal structures, there was a good representation of all four classes (Figure 3a). All but two of the structures were from lower organisms (virus, bacteria, archaebacteria, tetrahymena), and these two structures were classed as TBM, which reflects the high coverage of fold space that has now been achieved in higher organisms (Table S1).

**Figure 3.**
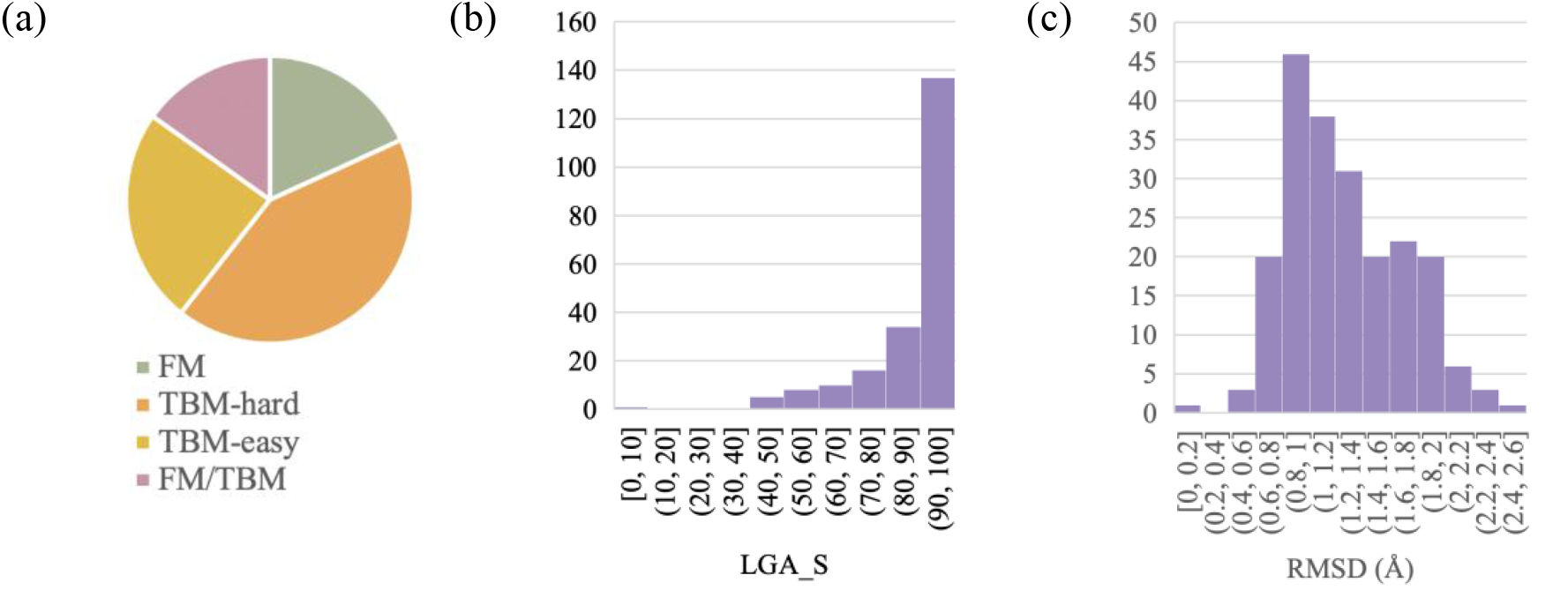
Classifications and accuracy for 34 crystal structures in CASP14. (a) Proportion of different modelling categories; FM (Free Modelling) TBM (Template Based Modelling). PDB entry 6vr4 was counted as a single FM target. Histogram of distribution of all 5 submitted AlphaFold2 models for 44 crystallographic targets of interest for (b) LGA_S and (c) RMSD.

### 3.4. Metrics for target quality

For the purposes of judging models for molecular replacement, the model *rmsd* and *f_m_* are the important metrics.

Of all the metrics reported by CASP, the sequence independent LGA(4Å) parameters RMSD and LGA_S are most closely allied with the *rmsd* and *f_m_*. The RMSD is the root-mean-square deviation for the subset of Cα atoms from the model that correspond to the residues from the target structure in the sequence-independent LGA superposition. The LGA_S is the sequence similarity score, which is a combination of the GDT score and the LCS score, where the LCS is the longest continuous segment (as a percentage of the total sequence) that can fit under a *rmsd* of a given cut-off. The LGA_S scores are similar to the GDT_TS scores for closely aligned structures. LGA_S is not sensitive to out of sequence register errors, in the same way that the *f_m_* of a model for molecular replacement phasing is not directly sensitive to any registration errors of the model.

A histogram of the metrics RMSD and LGA_S for the AlphaFold2 models across the 44 CASP crystallographic targets of interest is shown in Figure 3(b,c). The LGA_S is skewed towards almost full sequence coverage with an average of 87% and the RMSD is clustered around the average RMSD of 1.27 Å.

The RMSD and LGA_S also show the superiority of the AlphaFold2 models over models submitted by other groups. Table 2 shows the metrics for the best LGA_S and best RMSD AlphaFold2 models (of the 5 submitted) for the 44 CASP crystallographic targets of interest, compared with the best models by the same metrics overall. In only two cases (T1073 and T1085) was an AlphaFold2 model not the best as scored by LGA_S. In the case of T1085, the difference in LCS_S was negligible (less than half a percent), and the AlphaFold2 model had a much lower RMSD (0.85 Å versus 1.39 Å). In the case of T1073, the differences between the models were mostly confined to a short region of N-terminal helix that extended from the body of the globular fold. In 15 cases a non-AlphaFold2 model had a lower RMSD, however this was exclusively at the expense of a lower (usually a far lower) LGA_S.

**Table 2.** Best models for the targets from 34 crystal structures included in CASP14. Model is given as (CASP group number)_(ranked model number). In brackets are LGA_S and RMSD.

| Crystal Number | Target(s) in ASU | Best AlphaFold2 by LGA_S | Best model overall by LGA_S | Best AlphaFold2 by RMSD | Best model overall by RMSD |
| --- | --- | --- | --- | --- | --- |
| 1 | T1024 | 427_3 [87.5, 1.83] | 427_3 | 427_1 [58.8, 1.60] | 427_1 |
| 2 | T1030 | 427_2 [62.0, 1.82] | 427_2 | 427_2 | 013_2 [39.2, 1.27] |
| 3 | T1031 | 427_2 [94.0, 1.12] | 427_2 | 427_4 [93.7, 0.98] | 427_4 |
|  | T1033 | 427_1 [93.3, 1.39] | 427_1 | 427_3 [92.5, 1.36] | 259_4 [39.3, 1.29] |
|  | T1035 | 427_5 [99.0, 0.81] | 427_5 | 427_5 | 427_5 |
|  | T1037 | 427_4 [95.4, 1.12] | 427_4 | 427_5 [93.7, 1.11] | 427_5 |
|  | T1039 | 427_1 [86.3, 1.61] | 427_1 | 427_1 | 071_1 [33.5, 1.17] |
|  | T1040 | 427_1 [77.5, 1.95] | 427_1 | 427_2 [76.3, 1.90] | 140_1 [16.4, 1.31] |
|  | T1041 | 427_1 [94.7, 1.21] | 427_1 | 427_1 | 427_1 |
|  | T1042 | 427_3 [93.8, 1.22] | 427_3 | 427_5 [93.4, 1.21] | 427_5 |
|  | T1043 | 427_3 [90.2, 1.42] | 427_3 | 427_1 [90.0, 1.41] | 427_1 |
| 4 | T1032 | 427_3 [71.1, 1.67] | 427_3 | 427_1 [70.1, 1.65] | 427_1 |
| 5 | T1034 | 427_1 [96.9, 1.00] | 427_1 | 427_2 [95.7, 0.87] | 427_2 |
| 6 | T1038 | 427_2 [91.9, 1.17] | 427_2 | 427_2 | 427_2 |
| 7 | T1046s1 | 427_4 [98.1, 0.68] | 427_4 | 427_1 [98.1, 0.64] | 427_1 |
|  | T1046s2 | 427_1 [98.9, 0.69] | 427_1 | 427_1 | 427_1 |
| 8 | T1048* |  |  |  |  |
| 9 | T1049 | 427_1 [95.3, 0.82] | 427_1 | 427_1 | 427_1 |
| 10 | T1050 | 427_1 [93.3, 1.26] | 427_1 | 427_1 | 427_1 |
| 11 | T1052 | 427_4 [63.4, 1.17] | 427_4 | 427_5 [62.9, 1.14] | 337_5 [45.5, 1.13] |
| 12 | T1053 | 427_3 [96.9, 0.98] | 427_3 | 427_3 | 427_3 |
| 13 | T1054 | 427_3 [93.7, 0.84] | 427_3 | 427_2 [93.0, 0.81] | 029_1 [49.7, 0.76] |
| 14 | T1056 | 427_2 [99.3, 0.66] | 427_2 | 427_2 | 427_2 |
| 15 | T1058 | 427_3 [93.7, 1.25] | 427_3 | 427_3 | 427_3 |
| 16 | T1064 | 427_1 [92.6, 1.34] | 427_1 | 427_2 [91.0, 1.31] | 427_2 |
| 17 | T1065s1 | 427_2 [98.4, 0.91] | 427_2 | 427_4 [97.8, 0.85] | 427_4 |
|  | T1065s2 | 427_1 [99.5, 0.60] | 427_1 | 427_1 | 427_1 |
| 18 | T1067 | 427_3 [92.9, 0.86] | 427_3 | 427_3 | 427_3 |
| 19 | T1070 | 427_5 [45.0, 1.69] | 427_3 | 427_3 [41.2, 1.52] | 334_1 [30.4, 1.22] |
| 20 | T1073 | 427_3 [86.7, 1.76] | 288_4 [95.2, 1.47] | 427_5 [85.5, 1.41] | 217_3 [83.8, 0.97] |
| 21 | T1074 | 427_2 [93.7, 1.15] | 427_2 | 427_4 [92.6, 1.06] | 427_4 |
| 22 | T1079 | 427_4 [96.7, 1.05] | 427_4 | 427_2 [96.7, 1.03] | 427_2 |
| 23 | T1080 | 427_4 [91.6, 1.37] | 427_4 | 427_4 | 427_4 |
| 24 | T1082 | 427_1 [97.9, 0.88] | 427_1 | 427_2 [97.3, 0.88] | 427_2 |
| 25 | T1083 | 427_4 [91.9, 1.09] | 427_4 | 427_4 | 392_2 [88.7, 0.96] |
| 26 | T1084 | 427_5 [94.6, 0.85] | 129_3 [95.1, 1.39] | 427_4 [94.0, 0.77] | 480_4 [92.6, 0.60] |
| 27 | T1085 | 427_1 [88.2, 1.86] | 427_1 | 427_1 | 473_4 [32.5, 1.45] |
| 28 | T1086 | 427_1 [89.6, 1.78] | 427_1 | 427_4 [88.7, 1.64] | 080_1 [53.0, 1.60] |
| 29 | T1087 | 427_3 [97.0, 0.63] | 427_3 | 427_2 [96.8, 0.57] | 081_3 [40.3, 0.38] |
| 30 | T1089 | 427_2 [99.0, 0.71] | 427_2 | 427_2 | 427_2 |
| 31 | T1090 | 427_3 [95.4, 1.16] | 427_3 | 427_1 [92.1, 1.04] | 427_1 |
| 32 | T1091 | 427_2 [79.8, 2.00] | 427_2 | 427_5 [77.0, 1.96] | 403_2 [27.3, 1.47] |
| 33 | T1100 | 427_2 [90.3, 1.68] | 427_2 | 427_4 [55.9, 1.06] | 132_2 [18.2, 0.81] |
| 34 | T1101 | 427_4 [92.0, 1.17] | 427_4 | 427_3 [91.9, 1.15] | 427_3 |

\* cancelled
Group numbers: 427 (AlphaFold2), 013 (FEIG-S), 029 (Venclovas), 071 (Kiharalab), 080 (FOLDYNE), 081 (MUFOLD), 129 (Zhang), 132 (PBuild), 140 (Yang-Server), 217 (CAO-QA1), 259 (AWSEM-CHEN), 288 (DATE), 334 (FEIG-R3), 337 (CATHER), 392 (trfold), 403 (BAKER-experimental), 473 (BAKER) and 480 (FEIG-R2).

These two metrics (LGA_S and RMSD from the LGA(4Å)) are not ideal metrics for representing *rmsd* and *f_m_*. Calculation of *rmsd* and *f_m_* is critically dependent on the alignment of the structures, and alignment should, properly, be based on electron density rather than coordinates, a problem that will be addressed elsewhere.

## 4. Molecular replacement methods

The high LGA_S and low RMSD scores of the best AlphaFold2 models indicated that AlphaFold2 models were good prospects for achieving phasing by molecular replacement.

Initially, the 31 targets with a one-to-one correspondence between a sequence and a CASP crystal structure, the 2 targets each for the 2 heterodimeric structures and the 9 targets for crystal 6vr4 were used for molecular replacement (in total, 44 CASP crystallographic targets to be placed in 34 crystal structures).

If molecular replacement with the target representing the full sequence failed, and the target was one of the 11 classified as ‘multidom’, then molecular replacement was attempted with the domains.

The AlphaFold2 models for any given target almost exclusively superimposed with very little coordinate variability and so creating an ensemble structure did not indicate where poorly modelled regions could be trimmed (by divergence between models) unless exceptionally small divergence distance thresholds were used (e.g. 0.1 Å). Rather than using a tiny deviation threshold, trimming was performed using a threshold for the predicted error per residue supplied as part of the AlphaFold2 structure prediction.

The AlphaFold2 models (full and domain targets, untrimmed and trimmed) were used for molecular replacement in *Phaser.voyager* (manuscript in preparation). *Phaser.voyager* uses the *phasertng* codebase (McCoy *et al*., 2020). The initial VRMS (effective *rmsd)* was set to 1.2 Å and then refined for posed models. The 5 submitted AlphaFold2 models were used as an ensemble. If the target structure was available, the pose was checked to see if it was a match for the target coordinates with *phenix.famos* (Caballero *et al*., 2018). To confirm the solution we used *phenix.autobuild*for initial R-value and R-free (Terwilliger *et al*., 2007). Structures were considered solved if the density correlation was under 0.3. If the R-free was high, model improvement was attempted with *phenix.morph_model*. Further manual model building and refinement was not pursued.

In one case (described below) molecular replacement with *Phaser.voyager* failed, and molecular replacement was performed with *Arcimboldo-lite* for coiled-coils (Caballero *et al*., 2018).

## 5. Molecular replacement results

Of the 34 crystal structures, 31 could be solved by molecular replacement with the AlphaFold2 models, 2 partially solved, and one could not be solved with the AlphaFold2 model.

The case that could not be solved with the AlphaFold2 model was crystal 8, the coiled-coiled structure 6un9, target T1048. Although not solvable with the full AlphaFold2 model, this structure could be solved with a generic 20-residue poly-alanine helix using *Arcimboldo_lite* for coiled coils (Caballero *et al*., 2018).

A partial solution was achieved for crystal 3, the polymerase structure 6vr4, for which 6 of the 9 constituent CASP targets (in two copies each) could be placed. The full structure was not designated as a CASP target; we are unable to ascertain whether the whole 2194 residue structure is tractable for AlphaFold2 prediction. Should the whole structure have been a target, and had an AlphaFold2 model been available, it is possible that such a model would have also succeeded in molecular replacement.

A partial solution was also achieved for crystal 2, the all-helical structure 6poo. The full structure was designated as a ‘multidom’ CASP target with two domains. The second domain T1030-D1 could be placed unambiguously. The first domain T1030-D2 could be placed by molecular replacement but gave a very high final R-free.

Of the 31 solved structures, 28 were solved straightforwardly with the default *Phaser.voyager* protocol (Table 3).

**Table 3.** Summary of phasing of the 34 crystal structures of interest included in CASP14 with AlphaFold2 models. Crystals listed in bold are discussed in the text.

| # | Target | PDB | dmin | Number of reflections | Z | Domains | Filter | TFZ | R-free | R-free after morphing |
| --- | --- | --- | --- | --- | --- | --- | --- | --- | --- | --- |
| 1 | T1024 | 6t1z | 2.9 | 12686 | 1 | 1 | No | 24.7 | 0.52 | 0.42 |
| 2† | **T1030** | **6poo** | **3.0** | **7525** | **1** | **2** | **No** | **20.4** | **0.53** |  |
| 3† | **T1031** | 6vr4 | 3.5 | 92907 | 2 |  |  |  |  |  |
|  | **T1033** |  |  |  |  |  |  |  |  |  |
|  | **T1035** |  |  |  |  |  |  |  |  |  |
|  | **T1037** |  |  |  |  |  |  |  |  |  |
|  | **T1039** |  |  |  |  |  |  |  |  |  |
|  | **T1040** |  |  |  |  |  |  |  |  |  |
|  | **T1041** |  |  |  |  |  |  |  |  |  |
|  | **T1042** |  |  |  |  |  |  |  |  |  |
|  | **T1043** |  |  |  |  |  |  |  |  |  |
| 4 | **T1032** | **6n64** | **3.3** | **27936** | **6** | **1** | **0.7** | **16.1§** | **0.47** | **0.44** |
| 5 | T1034 | 6tmm | 2.1 | 47702 | 4 | 1 | No | 31.9 | 0.45 |  |
| 6 | T1038 | 6ya2 | 2.5 | 20426 | 3 | 1 | No | 24.3 | 0.35 |  |
| 7 | T1046s1 | 6px4 | 1.7 | 69112 | 2 | 2 | No | 31.8 | 0.35 |  |
|  | T1046s2 |  |  |  |  |  |  |  |  |  |
| 8‡ | **T1048\*** | **6un9** | **2.8** | **19203** | **4** |  |  |  |  |  |
| 9 | T1049 | 6y4f | 1.8 | 12228 | 1 | 1 | No | 23.7 | 0.34 |  |
| 10 | T1050 |  | 2.7 | 97731 | 3 | 1 | No | 39.0 | 0.30 |  |
| 11 | T1052 |  | 2.0 | 88914 | 2 | 1 | No | 48.6 | 0.43 |  |
| 12 | T1053 |  | 3.2 | 49627 | 4 | 1 | No | 51.0 | 0.35 |  |
| 13 | T1054 |  | 1.7 | 25547 | 1 | 1 | No | 33.2 | 0.34 |  |
| 14 | T1056 | 6yj1 | 2.3 | 17863 | 2 | 1 | No | 19.7 | 0.37 |  |
| 15 | T1058 |  | 3.1 | 20228 | 2 | 1 | No | 25.5 | 0.44 |  |
| 16 | T1064 | 7jtl | 2.0 | 16787 | 2 | 1 | No | 19.1 | 0.41 |  |
| 17 | T1065s1 |  | 1.6 | 35695 | 1 | 2 | No | 48.7 | 0.22 |  |
|  | T1065s2 |  |  |  |  |  |  |  |  |  |
| 18 | T1067 |  | 1.4 | 51025 | 1 | 1 | No | 57.2 | 0.26 |  |
| 19 | T1070 |  | 2.5 | 25412 | 1 | 1 | No | 6.8 | 0.49 | 0.42 |
| 20 | **T1073** |  | **1.9** | **27326** | **4** | **1** | **1** | **24.1** | **0.39** |  |
| 21 | T1074 |  | 1.5 | 25800 | 1 | 1 | No | 21.1 | 0.29 |  |
| 22 | T1079 |  | 3.2 | 47985 | 4 | 1 | No | 33.6 | 0.38 |  |
| 23 | **T1080** |  | **1.7** | **100570** | **6** | **1** | **No** | **20.9** | **0.39** |  |
| 24 | T1082 | 6x6o | 1.1 | 97672 | 2 | 1 | No | 33.6 | 0.44 | 0.31 |
| 25 | T1083 |  | 1.3 | 81236 | 4 | 1 | 0.6 | 30.0 | 0.48 | 0.35 |
| 26 | T1084 |  | 1.9 | 23901 | 3 | 1 | No | 32.1 | 0.38 |  |
| 27 | T1085 |  | 2.5 | 10758 | 1 | 3 | No | 22.0 | 0.40 |  |
| 28 | T1086 |  | 2.3 | 21887 | 1 | 1 | No | 18.7 | 0.41 |  |
| 29 | T1087 |  | 1.4 | 69617 | 4 | 1 | No | 43.2 | 0.25 |  |
| 30 | T1089 |  | 2.2 | 55192 | 2 | 1 | No | 63.5 | 0.29 |  |
| 31 | T1090 | 7k7w | 1.8 | 22947 | 1 | 1 | No | 27.2 | 0.29 |  |
| 32 | T1091 |  | 2.2 | 62789 | 1 | 4 | No | 22.5 | 0.44 |  |
| 33 | **T1100** |  | **3.1** | **36829** | **4** | **1** | **No** | **31.1** | **0.48** | **0.45** |
| 34 | T1101 |  | 1.4 | 58030 | 1 | 1 | No | 16.2 | 0.34 |  |

† partial solution
‡ no solution
§ placement of dimer

Details are given below for crystal 8 (no solution); crystals 2 and 3 (partial solutions); and crystals 20, 23 and 33 for which the structure solution was successful but proved more problematic.

### 5.1. Crystal 2

T1030 (PDB 6poo) is a helical bundle classified as ‘multidom’ with two domains. D2 could be placed unambiguously by molecular replacement, but the best pose for D1 was only able to superimpose a portion of the fragment, and the R-free after molecular replacement was greater than 0.50.

The overall C*α* rmsd of the first ranked AlphaFold2 model to the target for D1 was 2.8 Å over 154 residues and for D2 was 1.2 Å over 119 residues.

The high R-free of the molecular replacement solution can be attributed to model/target differences in D1 bending angles and angular disposition of the 6 constituent helices. Since the helices are long (residue lengths of 18, 15, 35, 15, 38 and 22) these differences result in systematic deviation of the coordinates, so that an overall *rmsd* does not give a complete picture of coordinate divergence.

Analysis with the HELANAL-PLUS server (Kumar & Bansal, 2012) showed that 6 AlphaFold2 helices were classified as [‘linear, ‘curved’, ‘linear, ‘linear, ‘unassigned, ‘curved’] while the target helices were classified as [‘curved’, ‘curved’, ‘kinked, ‘linear, ‘kinked, ‘curved’] based on average and maximum bending angles. Analysis with CCP4 helixang (Winn *et al*., 2011) gave the angles between helical axes of helix 1 and [2-6] for the AlphaFold2 model of [173°, 7°, 194°, 19°, −154°] and for the target of [172°, 5°, 171°, −23°, and −153°]; most notable were the differences in the disposition of helices 1-4 (Δ 23°) and 1-5 (Δ 42°).

### 5.2. Crystal 3

For CASP14, the single polypeptide chain of the virion-packaged DNA-dependent RNA polymerase of crAss-like phage phi14:2 was divided into 9 assessment domains (T1031 T1033 T1035 T1037 T1039 T1040 T1041 T1042 T1043), which we here refer to by numbering 1-9 (Figure 4). Eight domains were classed as FM, and one as FM/TBM.

**Figure 4.**
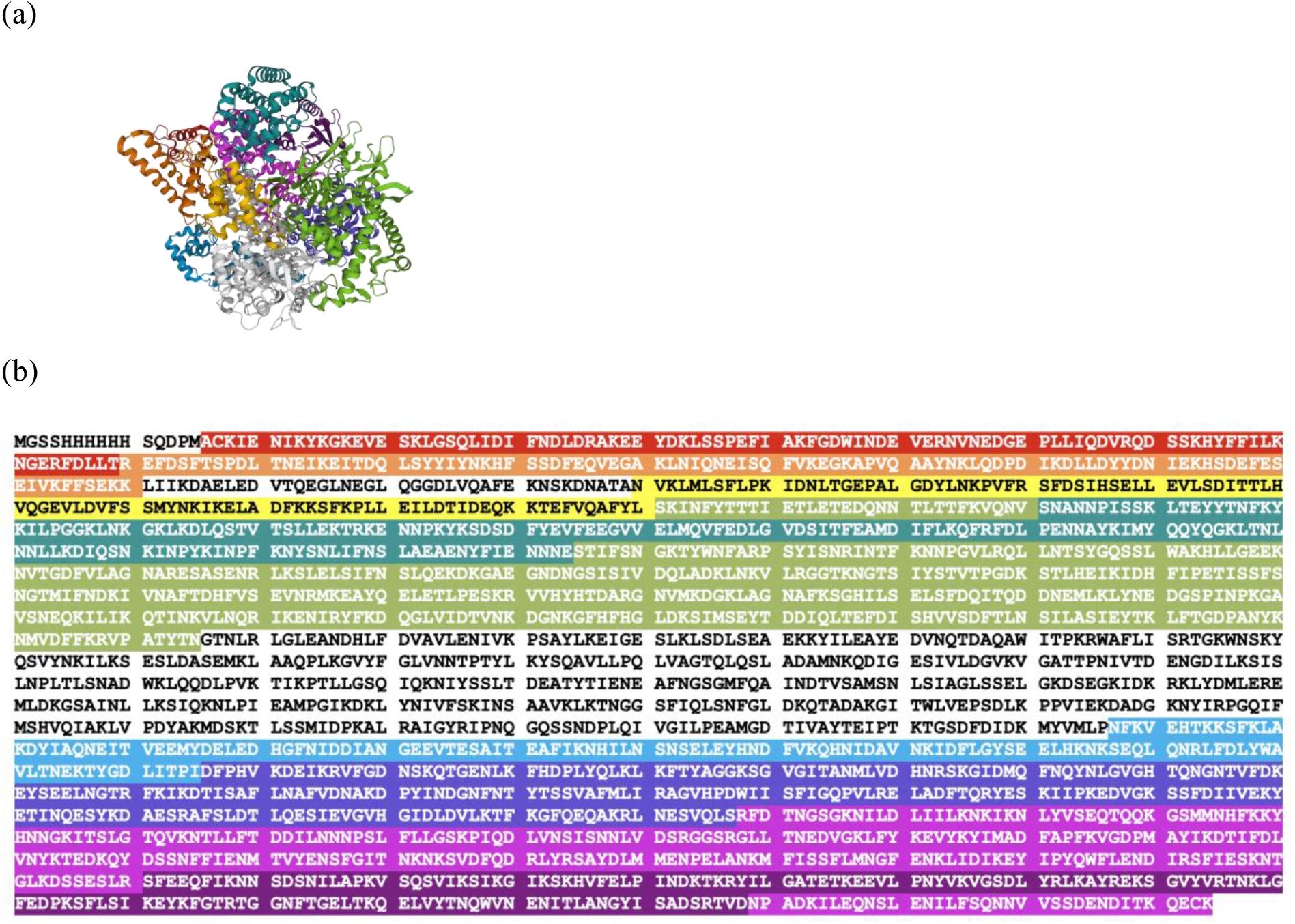
Crystal 3, PDB 6vr4, targets T1031 (1: red), T1033 (2: orange), T1035 (3: yellow), T1037 (4: khaki), T1039 (5: green), T1040 (6: blue), T1041 (7: purple), T1042 (8: magenta), T1043 (9: violet). Two targets are discontinuous in the primary sequence. (a) Structure with targets highlighted; regions not corresponding with a target are shown in grey. Figure created with Mol* (Sehnal *et al*., 2021). (b) Sequence with targets highlighted, regions not highlighted were not included in targets

There were two copies of the monomer in the asymmetric unit (PDB 6vr4; (Drobysheva *et al*., 2021)), related by a non-crystallographic two-fold. The assessment domains were used as models for molecular replacement. In total, 12 of the 18 domains could be placed, giving a ⅔ complete solution, which was insufficient for phasing the remaining fragments of the structure given the limited resolution of 3.5 Å. The partial solution was achieved by running *Phaser* from the command line. Domains were placed sequentially and the order of placement for the 12 domains was [4,4,7,7,2,3,2,3,8,8,5,5]. The second copies of domains 2, 3 and 8 were not placed by molecular replacement, but by applying the non-crystallographic symmetry operator to the already placed copies and performing rigid body refinement. After the placement of the first domain 2, 40 cycles of Refmac (Murshudov *et al*., 2011) refinement were performed to improve the partial structure before continuing. This procedure was repeated after placing the second domain 3. Domains 1, 5 and 9 could not be placed; these domains had very high *rmsd* to the target, over 2.5 Å (Table 4).

**Table 4.** Phasing of crystal 3 (PDB 6vr4) with AlphaFold2 models. For CASP14, the single polypeptide chain of the target sequence was divided into 9 assessment domains. There were two copies of the target sequence in the crystallographic asymmetric unit. Two copies each of six targets were found by molecular replacement and given chain identifiers A-L in order of placement. Targets T1031, T1040 and T1043 could not be placed.

|  | Model | Z | Type | residues | rmsd | Chain |
| --- | --- | --- | --- | --- | --- | --- |
| 1 | T1031TS427_1-D1 | 2 | FM | 95 | 2.91 |  |
| 2 | T1033TS427_1-D1 | 2 | FM | 100 | 1.58 | EG |
| 3 | T1035TS427_1-D1 | 2 | FM/TBM | 102 | 0.81 | FH |
| 4 | T1037TS427_1-D1 | 2 | FM | 404 | 1.25 | AB |
| 5 | T1039TS427_1-D1 | 2 | FM | 161 | 2.50 | KL |
| 6 | T1040TS427_1-D1 | 2 | FM | 130 | 2.76 |  |
| 7 | T1041TS427_1-D1 | 2 | FM | 242 | 1.70 | CD |
| 8 | T1042TS427_1-D1 | 2 | FM | 276 | 1.79 | IJ |
| 9 | T1043TS427_1-D1 | 2 | FM | 148 | 2.46 |  |

### 5.3. Crystal 4

T1032 was classified as FM/TBM. There were 6 copies of the sequence in the asymmetric unit (PDB 6n64; (Chen *et al*., 2020)) in 3 dimers.

The model’s LGA_S was 70%, with an RMSD of 1.7 Å. Successful molecular replacement required finding the portion of the model that was correct. The structure could be solved with two different approaches.

The first approach used *Arcimboldo_shredder* (Millán *et al*., 2018), which ‘shreds’ the model into fragments defined by spheres around each C*α* and uses the persistence of solutions across searches with different fragments as a way of enhancing molecular replacement signal. Four of the six copies of the structure in the asymmetric unit were initially found using the molecular replacement protocol described above. These formed two dimers, each with a non-crystallographic two-fold. One dimer was extracted from the partial solution and used successfully for molecular replacement to place the final two components. Extracting oligomeric associations from a partial structure solution and using them for completing the asymmetric unit is an established protocol in molecular replacement.

The second approach used the *Phaser.voyager* pipeline, after trimming the model where the predicted deviation between model and target (as generated by AlphaFold2) was greater than 0.7 Å. After successful molecular replacement to place all six copies with the protocol described, the complete AlphaFold2 model was superimposed on the fragment used for molecular replacement, and refined with *Phenix.morph_model* (which applies smooth distortions) to bring the R-free to 42%.

### 5.4. Crystal 8

T1048 (PDB 6un9) is a single helix and was cancelled from CASP14 on 20-10-2020 for ‘lack of tertiary structure’. However, a model for this structure was generated by AlphaFold2 before the target was cancelled.

This structure is a 61 residue coiled-coil. Coiled-coil structures are notoriously difficult to solve by molecular replacement because of the modulation of the data due to the helical repeats (Caballero *et al*., 2018; Thomas *et al*., 2015).

The structure had four copies of the target in the asymmetric unit. The overall C*α* rmsd of the AlphaFold2 model to chain A in the target structure was 2.1 Å for 67 residues. Analysis with HELANALPLUS server (Kumar & Bansal, 2012) showed that the maximum bending angle was markedly different (7.4° versus 25.7°); the model helix was classified as ‘curved’ while the target was classified as ‘kinked’. The structure could not be solved with the AlphaFold2 model in *Phaser.voyager*, even with the model optimized by trimming to reduce the *rmsd* to the deposited structure (at the expense of a lower fraction scattering).

*Arcimboldo_lite* in the coiled-coil mode was able to solve the structure using a generic 20-residue poly-alanine helix. The advantage of using short generic helices for coiled coil structures is that they are able to superimpose with multiple short sections of the coiled-coil helices with low *rmsd*. Structure solution required the ‘verification’ step, which is a powerful method for distinguishing the true solution from the abundance of false solutions that arise merely from helical placements that satisfy the helical modulations of the data (Caballero *et al*., 2018).

### 5.5. Crystal 20

All groups modelled the 12 residue N-terminal helix of T1073 with high *rmsd* to the target. This helix extends from the compact body of the fold. Structure solution was achieved by removing all sections of the Alphafold2 model with a predicted error over 1 Å, in a standard model preparation protocol with *Phaser.voyager*.

The challenge in this case was with data preparation not model preparation; the model preparation was unremarkable, but we found that this case also required some additional attention be given to the crystallographic data. A number of different datasets were available in the file provided. Molecular replacement was achieved, after *phenix.xtriage* (Zwart et al., 2005) analysis, with one of the datasets and with the resolution restricted to 2.8 Å.

### 5.6. Crystal 23

T1080 was classified as FM/TBM. There were 6 copies of the target in the asymmetric unit in two trimers.

This was the only case where the five submitted AlphaFold2 models showed significant deviation. Model 3 differed from the consensus fold of the other 4 in the first 40 N-terminal residues; these 40 residues took a very different conformation in model 3. In analysis, the consensus fold of 4 was correct, and model 3 incorrect, although the incorrect conformation could be considered as a “trimer swap” error, with the chain partly following the fold of a neighbouring monomer in the trimer. The molecular replacement model trimmed these residues and residues with a predicted deviation of more than 1.2 Å, leaving 78 of 133 residues. The molecular replacement model was therefore 60% of the target structure. After structure solution, the full AlphaFold2 consensus fold was superimposed on the solution and used for refinement.

### 5.7. Crystal 33

T1100 was classified as ‘multidom’ with two domains. There were four copies in the asymmetric unit in two dimers with translational non-crystallographic symmetry between the two dimers. D2 is a compact globular structure. D1 is a helical bundle structure with 4 helices of 52, 11, 64 and 28 residues. Within the dimer, the D1 helices formed a coiled-coil.

D2 could be placed unambiguously by molecular replacement.

D1 was more difficult to place. The problem can be attributed to model/target differences in helical bending angles. In general, the helices in the model were straighter than those in the target, with average bending angles of [4.5°, 7.1°, 4.6° and 3.6°] versus [8.5°, 9.3°, 8.5° and 7.8°] respectively. When the difference in bending angles were compounded over the long helices, particularly helix 1 (~75 Å) and 3 (~90 Å), it was not possible to simultaneously superimpose both ends of the model and target. Molecular replacement gave several closely related poses for D1, superimposing different portions of the model and target helices.

## 6. Survey of phasing methods

To discern the impact of high-accuracy *in silico* models on crystallographic phasing methods we undertook a survey of crystallographic phasing methods since the turn of the millennium.

We can divide crystallographic phasing strategies into four broad categories: direct methods, experimental phasing, molecular replacement, and difference Fourier methods (Fourier synthesis). The use of direct methods phasing is negligible for macromolecular crystallography, in contrast to its supreme dominance in small molecule crystallography (Sheldrick, 2008). Within the experimental phasing category are MAD (multi-wavelength anomalous dispersion), SAD (single-wavelength anomalous dispersion) and various IR (isomorphous replacement) methods; SIR (single isomorphous replacement), SIRAS (single isomorphous replacement with anomalous scattering) MIR (multiple isomorphous replacement), and MIRAS (multiple isomorphous replacement with anomalous scattering) (for review see (Rupp, 2010)).

The PDB mostly records macromolecular crystal structures that are published in peer-reviewed journals. The PDB is a record of novel crystal forms, if not novel structures. Our analysis only included those entries where protein was a component of the crystal.

The phasing method for each PDB entry is recorded in the ‘structure determination method’ field, which should allow a survey of phasing methods, however, analysis is not straightforward for a number of reasons listed below.

1. Although the ‘structure determination method’ field has been compulsory for submissions commenced after 29^th^ January 2019, a significant portion of the historical entries are null. Entries recorded before 2000 were regarded as too sparse for analysis. Null entries may be biased towards particular categories of phasing.
2. Although the ‘structure determination method’ field has been restricted to a few text strings for submissions commenced since 29^th^ January 2019, historically it was ‘free format’ and highly variable. For this study, all historical text entries were scanned by eye to assign each to one of the new restricted values. If the field referenced a number of methods (e.g. SAD with molecular replacement, or SIRAS/MAD) then the most senior phasing method was allocated with the order of precedence being miras, mir, mad, siras, sir, sad, molecular replacement and fourier synthesis.
3. Phasing by direct methods were not included in the study because a survey of entries with the ‘structure determination method’ field ab initio showed that, although these entries included those phased with direct methods, in the majority of cases ab initio referred to fragment-based approaches to molecular replacement or the use of direct methods for anomalous substructure determination. Since very few entries were categorised as ab initio, removing these from consideration did not significantly bias the results.
4. Checking a small sample of the entries in the ‘structure determination method’ field against the method recorded in the corresponding publication showed that the field was not always accurate. Inaccuracies may be biased towards particular categories of phasing.
5. Each entry has a deposition date, a release date and a revision date, therefore dating each entry is problematic. The deposition and release date are commonly separated by a year, but can be three years apart or even more. Revision dates are commonly very recent, as they include PDB-wide changes to PDB nomenclature. Only entries with “entity id” with value “1” were considered. In order to track the evolution of structure determination methods, we considered 5-year intervals, and restricted PDB identifiers for each interval such that both the deposition date *and* the release date were within the 5-year interval. Hence, our analysis sampled a subset of the PDB entries.
6. The understanding of the definition of different phasing methods may vary between crystallographers. For example, there is a degree of overlap between fourier synthesis and molecular replacement methods, as the former can be considered to be the latter but without an initial wide-radius search strategy (a search strategy employing rotation and translation functions); if the pose is outside the radius of convergence of rigid-body refinement, then (nominally) isomorphous crystals cannot be phased by difference Fourier methods and molecular replacement with a local or global search is used. Similarly, crystallographers may not distinguish between various similar experimental phasing methods (e.g. SAD versus SIRAS).

Despite these caveats, the trend in the change of structure determination methods over the last 20 years is clear (Figure 5a) and mirrors anecdotal experience. Molecular replacement now accounts for around 80% of phasing, increasing from around 50% of phasing in 2000, and molecular replacement and Fourier synthesis (difference Fourier) methods combined account for 95% of phasing. It is possible that an even higher proportion of structures were amenable to phasing by molecular replacement, had it been attempted.

**Figure 5.**
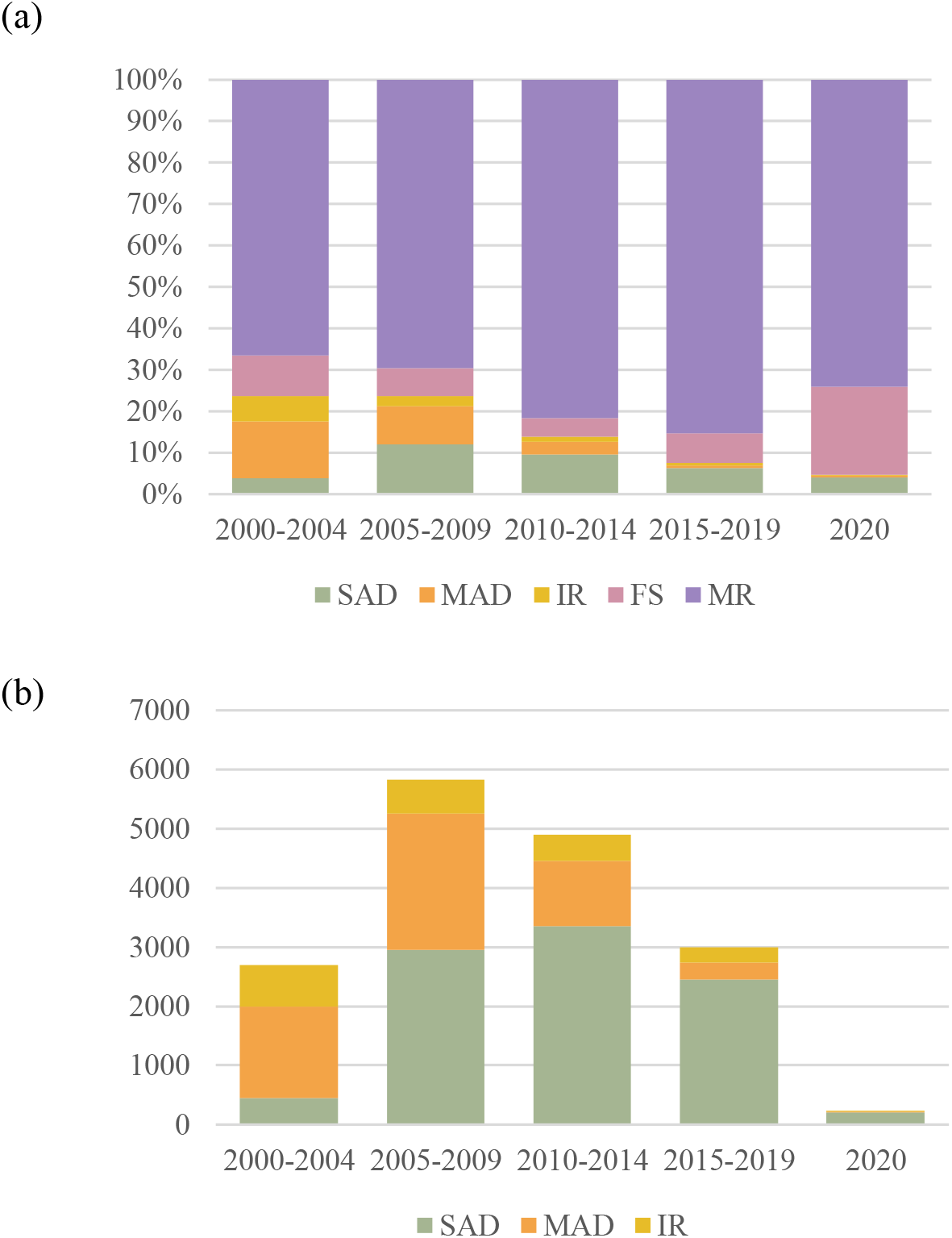
Phasing methods since 2000 as recorded in the ‘structure determination method’ field in the PDB; SAD (Single-wavelength Anomalous Dispersion), MAD (Multi-wavelength Anomalous Dispersion), IR (Isomorphous Replacement), FS (Fourier Synthesis) and MR (Molecular Replacement). Both the deposition and release date for the submission were within the 5-year time period shown. (a) All methods as percentage of PDB submissions per time period. (b) Experimental phasing methods as number of PDB submissions per time period.

Within the experimental phasing strategy, the method of experimental phasing has changed considerably, with MAD dominating in 2000, but SAD dominating today. SAD phasing, commonly using seleno-methionine substituted protein, now accounts for 82% of experimentally phased structures.

Of particular note is the decline in IR methods, originally the backbone of macromolecular phasing. To get an overview of the type of structures currently requiring IR to phase, we examined the structures submitted and released in 2020 in more detail (Table S2). 13 structures met this criterion, were phased by IR methods, and had publications available at the time of writing. Of the 13, we found one was actually solved by Se-SAD, one by Os-SAD, one by Pt-MAD and one by MR. Of the 9 confirmed examples of IR, only two structures used multiple derivatives.

## 7. Discussion

Self-evidently, crystallography requires crystals; crystallization is a bottle-neck, albeit one that has become far less constraining with advances in expression systems, fluidics, robotics, and computer vision. Not only must there be crystals, the crystals must diffract to better than 4 Å resolution to be useful for structural biology. With the data collected to the highest resolution a crystal form (space group, unit cell, asymmetric unit contents) will allow, it is usual to regard the crystallographic data as the ‘fixed’ component of molecular replacement phasing and to regard the model as the ‘dynamic’ component. Phasing pipelines are primarily designed for the automated exploration of many model structures prepared in different ways, with the hope that one will be accurate enough to be placed and allow model building and refinement to proceed with the single dataset provided.

To some degree, the AlphaFold2 models upend this paradigm. The need for extensive generation of models through different combinations of homologs in ensembles, different levels of trimming, the mining of domain databases, and the use of small secondary structure elements as models, is likely to be greatly reduced as these high-accuracy models become available. In essence, the AlphaFold2 models distil the information from all these methods, and more, in a single structure. The crystallographic problem may become one of finding a crystal form (for example, fewer copies in the asymmetric unit) that makes it amenable to molecular replacement with the *in silico* model(s). As an example of this, successful molecular replacement with T1073 (crystal 20) was achieved after massage of the data, rather than the model.

For 5 of the 19 structures that were classified as ‘multidom’, we used the domains, rather than the model of the whole structure, for molecular replacement. In this approach, the whole structure is built up by addition as domains are placed sequentially in the asymmetric unit. It is necessary when the disposition of the domains in the target is largely determined by crystal packing or allosteric effects.

The structures that were the most challenging to solve with the AlphaFold2 models contained extended helices. The problem was two-fold. Firstly, although helical secondary structure is very amenable to prediction, the subtle bends and kinks in the helices are more elusive, and these have long-range effects in the fit of the model to the target. Secondly, coiled-coils induce modulations in the diffraction data that confound the maximum likelihood targets in molecular replacement, a known issue and an active area of crystallographic methods development.

The statistics in Table 3 show that molecular replacement with the AlphaFold2 models, followed by simple refinement strategies, does not give structures suitable for immediate submission to the PDB. Investment in manual model building, informed by a degree of biological understanding of the structures, would have been required to obtain final structures, which was beyond the scope of this study.

The use of *in silico* models for molecular replacement will also impact downstream model building and refinement. Model building and refinement can already be assisted by techniques borrowed from *ab initio* modelling (Terwilliger *et al*., 2012). With models representing 100% of the polypeptide chain in approximately the correct conformation, model building is directed towards local minimization into electron density rather than *de novo* model building. In this study, we used *phenix.morph_model* to improve parts of the structure with initially poor fit to the density. In regions where the electron density is weak or absent due to static disorder in the crystal, constraining the structure to the model may lower R-factors and improve interpretation of the density. *In extremis*, the diffraction data may not need to be as good as it would need to be to refine, pass validation metrics, and publish the structure in the absence of the model. In effect, the diffraction data need only verify the model.

There is some work to do to optimize the use of high accuracy *in silico* models for the purposes of molecular replacement. The lack of conformational variability in the models is different from models drawn from homologs. Whereas homologs tend to vary most in the regions where they also deviate from the target structure, the AlphaFold2 models are very consistent (insistent) in regions even where they differ from the target structure. If taken purely at face value, this will lead to, for example, rejection of molecular replacement solutions due to (false) packing clashes. We can also improve how we make use of the estimated error in the coordinates in model preparation. It is also likely that improvements can be made in the estimation of σ_A_ for these models, since optimization of σ_A_ estimation has been calibrated for homologs rather than *in silico* models (Hatti *et al*., 2020).

The CASP14 crystal structures mostly represent a particular type of crystal structure: those that have a single protein sequence in the asymmetric unit and consist of one or few domains where the domain is unrelated, or poorly related, to known structures. These types of crystal structures are selected for by CASP, since they represent the more challenging structures for structure prediction. However, in support of structural biology, crystallography often focusses on protein complexes with peptide motifs, oligomeric associations, and multi-domain structures, often with domains that already have homologous structures in the PDB. That these can already be solved by molecular replacement or Fourier synthesis in at least 95% of cases is evident in the statistics and does not diminish their scientific interest.

Our survey of phasing methods indicates that IR phasing is becoming a specialist method. Despite the undoubted power of IR to obtain spectacularly good phases, even with low resolution and poor data, there are other factors that mean IR is avoided where possible. If using heavy metals, it requires handling high toxic metal salts, which also bind to protein crystallographers, not just proteins (Blundell & Johnson, 1976). Methods that incorporate Noble gasses such as xenon require specially designed high-pressure cells and appropriate training and support to use. We note that 2 of the 9 structures phased by IR in our survey of 2020 were phased by SIRAS using iodine, which is a non-toxic and simple method.

Crystallographic phasing strategies have evolved continuously since 1913 (Ewald, 1962; Brooks-Bartlett & Garman, 2015), and the contribution of the high-accuracy models will continue this evolution. It will enable crystallographers to concentrate their efforts even more keenly on the structural biology by making crystallographic phasing even more straightforward. We should look forward to the biological insights that this will bring.

## Acknowledgements

We thank all contributors to CASP14, in particular the authors who gave access to their data ahead of deposition in the PDB. We also thank Rachel Kramer Green at RCSB for supplying information about PDB-entry record history. This research was supported by funding from a Wellcome Trust Principal Research Fellowship to RJR (grant 209407/Z/17/Z).

---

**Table S1.**
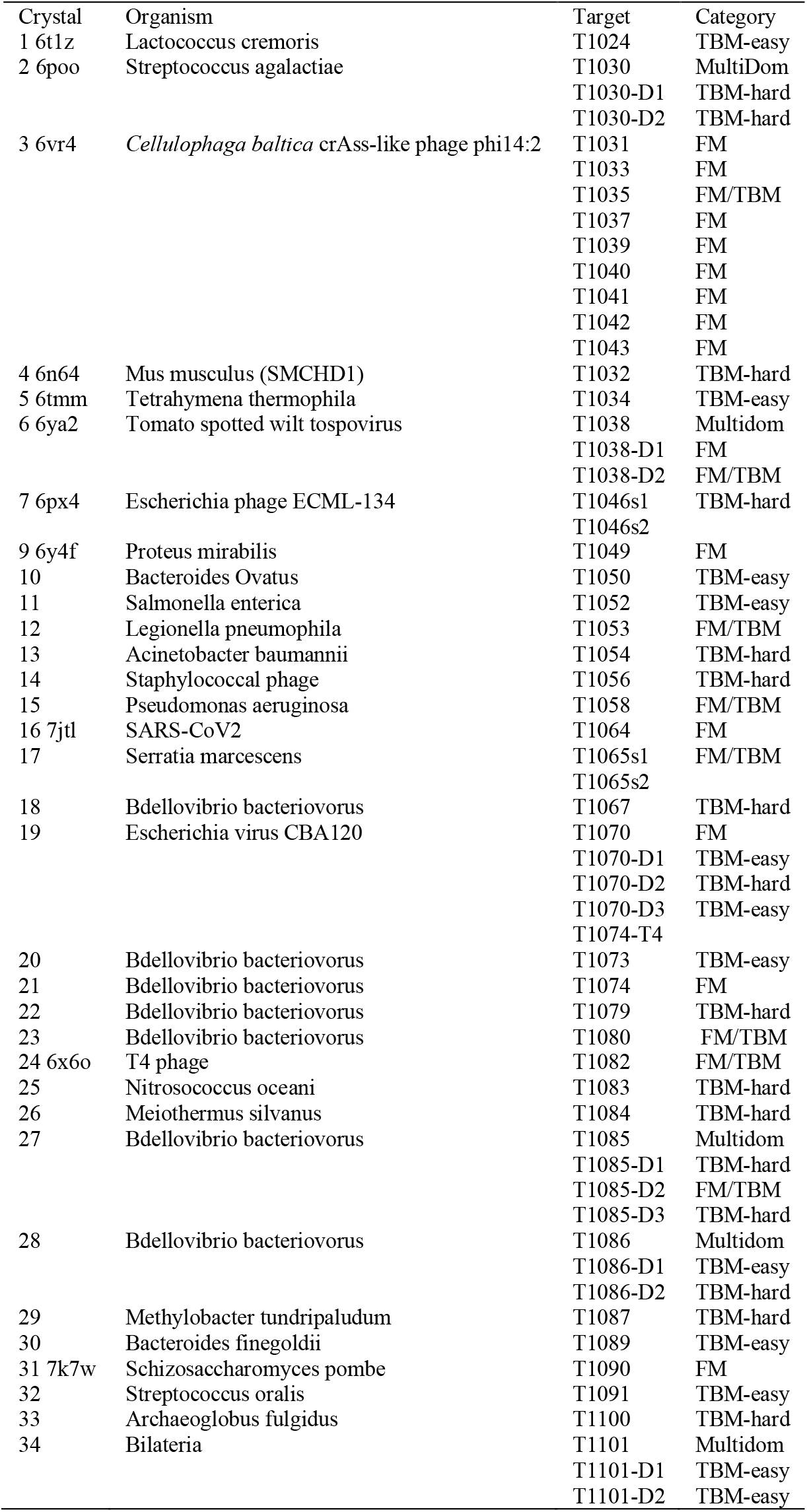
Crystallographic targets of interest showing organism of origin and prediction classification. Free modelling (FM), template-based modelling (TBM) and borderline between free modelling and template-based modelling (FM/TBM).

**Table S2.** Crystals phased by isomorphous replacement, that were reported to the PDB, contained protein and had both a deposition date and release date in 2020. PDB phasing method as recorded in the ‘structure determination method’ field.

|  | PDBID | resolution | PDB Phasing Method | Publication Phasing Method | Details | Publication |
| --- | --- | --- | --- | --- | --- | --- |
| 1 | 6TUG | 2.4 | SIRAS | SIRAS | native + Hg | (Garcia-Doval *et al.*, 2020) |
| 2 | 6VJI | 2.5 | MIRAS | MAD/MIRAS | Se-MAD + Pt + Au + native | (Eckenroth *et al.*, 2021) |
| 3 | 6VW9 | 1.8 | SIRAS | SIRAS | native + I | (Zimanyi *et al.*, 2020) |
| 4 | 6W8R | 2.8 | MIRAS | SIRAS | native + Se | (Zhu *et al.*, 2020) |
| 5 | 6WA6 | 2.8 | MIRAS | MIRAS | Au + Pt + U + native | (Jensen *et al.*, 2020) |
| 6 | 6WHB | 1.9 | SIRAS | SIRAS | native + I | (Filipčík *et al.*, 2020) |
| 7 | 6WN2 | 1.8 | MIR | MR |  | (He *et al.*, 2020) |
| 8 | 6Y7F | 2.1 | SIRAS | SIRAS | native + Hg | (Nie *et al.*, 2020) |
| 9 | 6Y9L | 4.1 | MIRAS | SAD | Os-SAD | (Bahat *et al.*, 2020) |
| 10 | 6YV7 | 2.7 | SIRAS | SIRAS | native + W | (Gandini *et al.*, 2020) |
| 11 | 6Z4E | 2.0 | MIR | MAD | Pt-MAD | (Micevski *et al.*, 2020) |
| 12 | 7AED | 1.8 | SIRAS | SAD | Se-SAD | (Jäger *et al.*, 2020) |
| 13 | 7BQI | 1.3 | SIRAS | SIRAS | native + Hg | (Sakurai *et al.*, 2020) |

